# Rapid and Efficient BAC Recombineering: Gain & Loss Screening System

**DOI:** 10.1101/2020.06.22.166017

**Authors:** Myeong Uk Kuk, Sekyung Oh, Joon Tae Park

**Affiliations:** Division of Life Sciences, College of Life Sciences and Bioengineering, Incheon National University, Incheon, Korea; Department of Medicine, Catholic Kwandong University College of Medicine, Incheon, Korea; Institute for Biomedical Research, Catholic Kwandong University International St. Mary’s Hospital, Incheon, Korea

**Keywords:** recombineering, lamda-red recombinase, gain & loss screening system

## Abstract

Recombineering has been developed to modify bacterial artificial chromosome (BAC) via homologous recombination. Nevertheless, as a screening strategy to identify the correct clone was not properly developed, it was difficult to obtain a correct clone within a limited time period. To address these issues, we developed a new screening method (a gain & loss screening system) that enables the efficient identification of the recombineered clone. Simple inoculation of cells into LB medium with appropriate antibiotics visually revealed the positive clones within 24 h. DNA sequencing confirmed 100% accuracy of this screening method by showing that all positive clones exhibited recombinant sequences. Furthermore, our new method allowed us to complete the entire procedure consisting of 1^st^ recombineering, flip-out and 2^nd^ recombineering in just 13 days. Overall, our new strategy may provide a new avenue for BAC recombeerining, as evidenced by markedly increased accuracy and subsequently shortened recombineering duration.

## INTRODUCTION

The production of biopharmaceutical proteins has been a key topic in biotechnology with mammalian cell culture being a widely used method for their production (Wurm 2004). Currently, over 100 therapeutic proteins are being produced in mammalian systems, and their number is expected to increase dramatically as new therapeutic antibodies are developed (Rita Costa et al. 2010). Thus, significant efforts have been made in the last decades to improve protein production in mammalian cell lines. Plasmid-based vectors are the most widely used tools for protein production. They include promoters that induce the expression of gene-of-interest (GOI). However, expression of GOI in plasmid-based vectors is greatly affected by surrounding chromatin at the integration site. Once the vector is integrated into a “silent chromatin” region, the expression tends to silence over time (i.e., positional chromatin effects) (Giraldo and Montoliu 2001). Accordingly, several strategies have been developed to avoid the local effects of chromatin. One of the most widely used methods is to use bacterial artificial chromosome (BAC) that can accommodate whole mammalian loci. BAC can accommodate a complete gene containing all *cis*-acting regulatory elements in their native configuration. Therefore, BAC is minimally affected by the surrounding chromatin at the integration site, so BAC can accurately deliver the expected expression pattern (Giraldo and Montoliu 2001).

Nevertheless, since the BAC size is very large, their modification cannot be made using standard cloning procedures (e.g., restriction enzyme digestion or ligation). BAC recombineering only allows the exchange of genetic information between two DNA molecules in an accurate, specific, and faithful way, regardless of the size of the DNA. However, BAC recombineering is very labor-intensive and time-consuming due to a large number of false positive background colonies during screening procedure. Therefore, BAC recombineering constituted a substantial barrier for less experienced researchers to consider BAC suitable as expression vectors for protein production.

Here we present a new screening method, the gain & loss screening system, which can provide markedly increased accuracy and shortened working time. This new strategy may provide powerful new tool for facilitating e.g. biopharmaceutical protein expression and other large-vector applications, and rendering such approaches feasible for less experienced laboratories.

## MATERIALS AND METHODS

### Bacterial strain and BAC clones

SW105 bacteria: Genotype *F-mcrA Δ(mrr-hsdRMS-mcrBC)* Φ*80dlacZ* M15 *ΔlacX74 deoR recA1 endA1 ara*D139 Δ(*ara, leu*) 7649 *galU galK rspL nupG* [*λcI857* (*cro-bioA*) <> *tet*] [(*cro-bio A*) <> *araC-PBADflpe*]. SW105 bacteria was generously provided by the Copeland laboratory at the National Cancer Institute. BAC clone (RP24-85L15) was purchased from BACPAC resources center (CHORI, Oakland, CA, USA).

### Preparation of BAC targeting cassette (BTC) and chloramphenicol (Cam) targeting cassette (CTC)

BAC targeting vector (BTV) (Addgene, cat. no. 131589) and Cam targeting cassette (CTV) (Addgene, cat. no. 131590) are available from Addgene. Incubate BTV with XhoI & XmaI and CTV with BamHI & XhoI for 4 h at 37°C. Load the digested DNA onto a 0.2% agarose gel. Run the gel for 30 min at 100 V on a DNA electrophoresis device (Bioand, cat. no. Mini-ES). Following electrophoresis, cut out the DNA band containing the BTC and CTC under a LED trans illuminator (Maestrogen, cat. no. SLB-01W). Purify the DNA using a gel extraction kit (Qiagen, cat. no. 28704). Measure the DNA concentration using Nanodrop spectrophotometer (Denovix, cat. no. DS-11) and dilute the DNA to 8–10 ng/μl using DW.

### Overview of the procedure

The comprehensive details on the methodology is describe in the Supplementary Material. To provide comprehensive details on the methodology, we have organized the BAC recombineering process into five consecutive stages, which are illustrated in a general flow-chart (Supplementary Fig. 1). In the first stage (steps 1–8), the identification of the desired BAC and the preparation of SW105 bacteria containing the BAC was performed. In the second stage (steps 9–15), the 1^st^ recombineering was conducted to introduce a BTC into the predetermined region of BAC. In the third stage (steps 16–18), flip-out was conducted to delete the selection marker, kanamycin resistance (*KanR*) gene. In the fourth stage (steps 19–23), the 2^nd^ recombineering was conducted to introduce a CTC. In the final stage (steps 24–27), maxi-preparation of BAC DNA was conducted.

## RESULTS

### 1^st^ gain & loss screening system

Two different BAC recombineering systems are widely used including those based on bacterial phage-encoded recombinases; one uses episomal plasmids to supply RecET of the Rac phage (Zhang et al. 1998; Muyrers et al. 2000) and the other utilizes a temperature-sensitive lamda repressor to control the expression of lamda-red recombinase (Yu et al. 2000; Lee et al. 2001). Lamda-red recombinase appears to be at least 50- to 100-fold more efficient than the RecET system (Zhang et al. 1996). Thus, methodology based on the lamda-red recombinase was selected in this study.

SW105 bacteria strain has been genetically modified to have the lamda-red recombinase system. It has a PL operon encoding lamda-red recombinase (*exo*, *bet*, and *gam*) that plays a crucial role in the recombineering process. The PL operon is under strict control of the temperature-sensitive lamda repressor (cI857). At low temperatures in the range of 30–34°C, cI857 is active and binds to the operator site thereby preventing transcription of the recombinant genes. Thermal upshift to 42°C reversibly inhibits the activity of cI857, thereby activating transcription of the recombinant genes.

For the recombineering, a 5’ homology region (HR) and 3’ HR was introduced into the BAC targeting vector (BTV, Addgene ID: 131589) containing GOI and a kanamycin resistant gene (*KanR*) (Fig. 1A). To purify BAC targeting cassette (BTC), the BTV with HRs was cut with appropriate XhoI and XmaI (Fig. 1A). As BTC contains *KanR* with two flanking FRT sites, the recombination of BTC with a BAC clone after brief heat shock at 42°C will result in a positive clone, which has dual antibiotic resistant genes, *KanR* (from BTC) and chloramphenicol resistant gene (*CamR*; from the BAC).

**Fig. 1.**
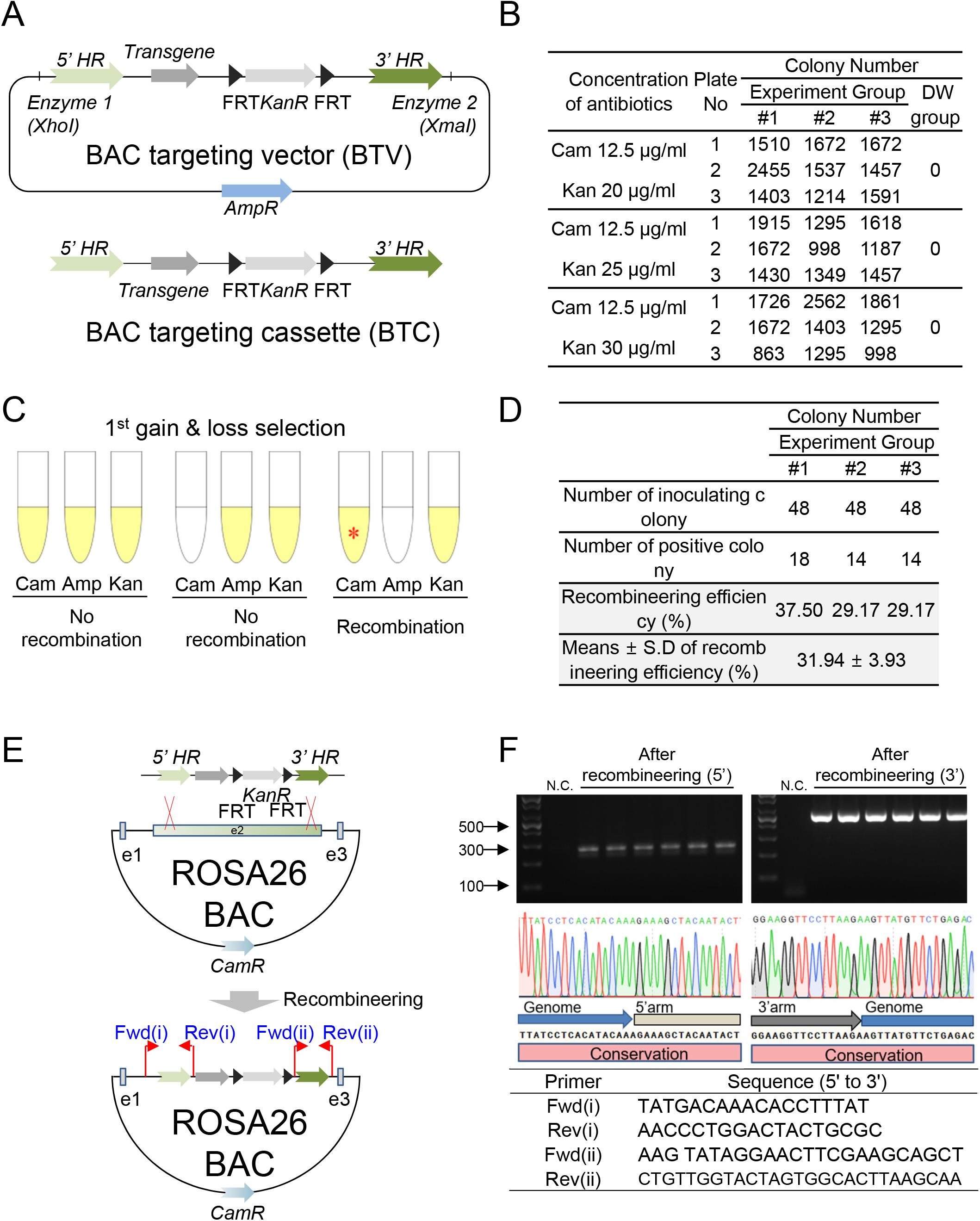
Detailed procedure and results for the 1^st^ recombineering. (A) Overview of the BAC targeting vector (BTV, Addgene ID: 131589) and BAC targeting cassette (BTC). (B) Summary of colony numbers in LB plates containing Cam and different concentrations of Kan following the 1^st^ BAC recombineering. (C) Illustration of the 1^st^ gain & loss selection system for identifying recombinant clones. Red asterisk (*) indicates the recombineering-positive clone. (D) Efficiency of the 1^st^ recombineering after performing the 1^st^ gain & loss selection system. Means ± S.D., N = 3. (E) 1^st^ recombineering scheme and position of colony PCR primers for recombination confirmation. (F) Picture of agarose gel electrophoresis showing colony PCR results. PCR product was only amplified in the recombineering-positive clone. Chromatogram results showing that the sequence of the amplified PCR product was conserved compared to the reference sequence at the genome-BTC boundary.

Following the recombineering, a LB plate containing Cam and Kan was used to discriminate the candidate clone(s). The number of candidate colonies appeared after 48 h incubation at 32°C was summarized in Fig. 1B. As BTV in supercoiled form cannot be efficiently cleaved with restriction enzymes, non-cleaved BTVs may be included in the purified BTC. BTV contamination will result in a non-recombinant clone, which has triple antibiotic resistant genes, an ampicillin resistant gene (*AmpR*; from BTV), *KanR* (from BTV) and *CamR* (from the BAC). To identify the clones derives from the desired recombination, the 1^st^ gain & loss screening system was applied (Fig. 1C). The candidate clones on the plate was inoculated in the 1^st^ gain & loss screening system and incubated for 24 h (Fig. 1C). This system will exhibit the three possible cases (Fig. 1C).

1. Case#1: positive in LB with Cam, positive with Amp, positive with Kan (no recombination)
2. Case#2: negative in LB with Cam, positive with Amp, positive with Kan (no recombination)
3. Case#3: positive in LB with Cam, negative with Amp, positive with Kan (recombination)

Case#3 only occurs if the recombineering is successful. The average efficiency of obtaining a case#3 was 31.94% (Fig. 1D). To confirm the correct recombination, colony PCR was conducted on all clones from case#3. The forward (Fwd) primer was located in the BAC vector and reverse (Rev) primer in BTC (Fig. 1E). The size of PCR product in 5’ recombineered and 3’ recombineered region was approximately 280 bp and 500 bp, respectively, indicating the successful recombineering (Fig. 1F). Following PCR, DNA sequencing was performed to verify the recombinant region. All positive clones were found to exhibit recombinant sequences indicating 100% accuracy of 1^st^ gain & loss screening system (Fig. 1F). Taken together, these results imply that simple inoculation of cells into the screening system visually displayed the positive clones with 100% accuracy.

### 2^nd^ gain & loss screening system

During the 1^st^ recombineering step, successful recombination was carried out through the introduction of *KanR*. However, *KanR* should be deleted for subsequent experiments. The SW105 strain harbors an endogenous L-arabinose-inducible *FLP* gene. Since *KanR* is flanked by two FRT sites, the induced FLP gene in the presence of L-arabinose will delete *KanR* (Fig. 2A).

**Fig. 2.**
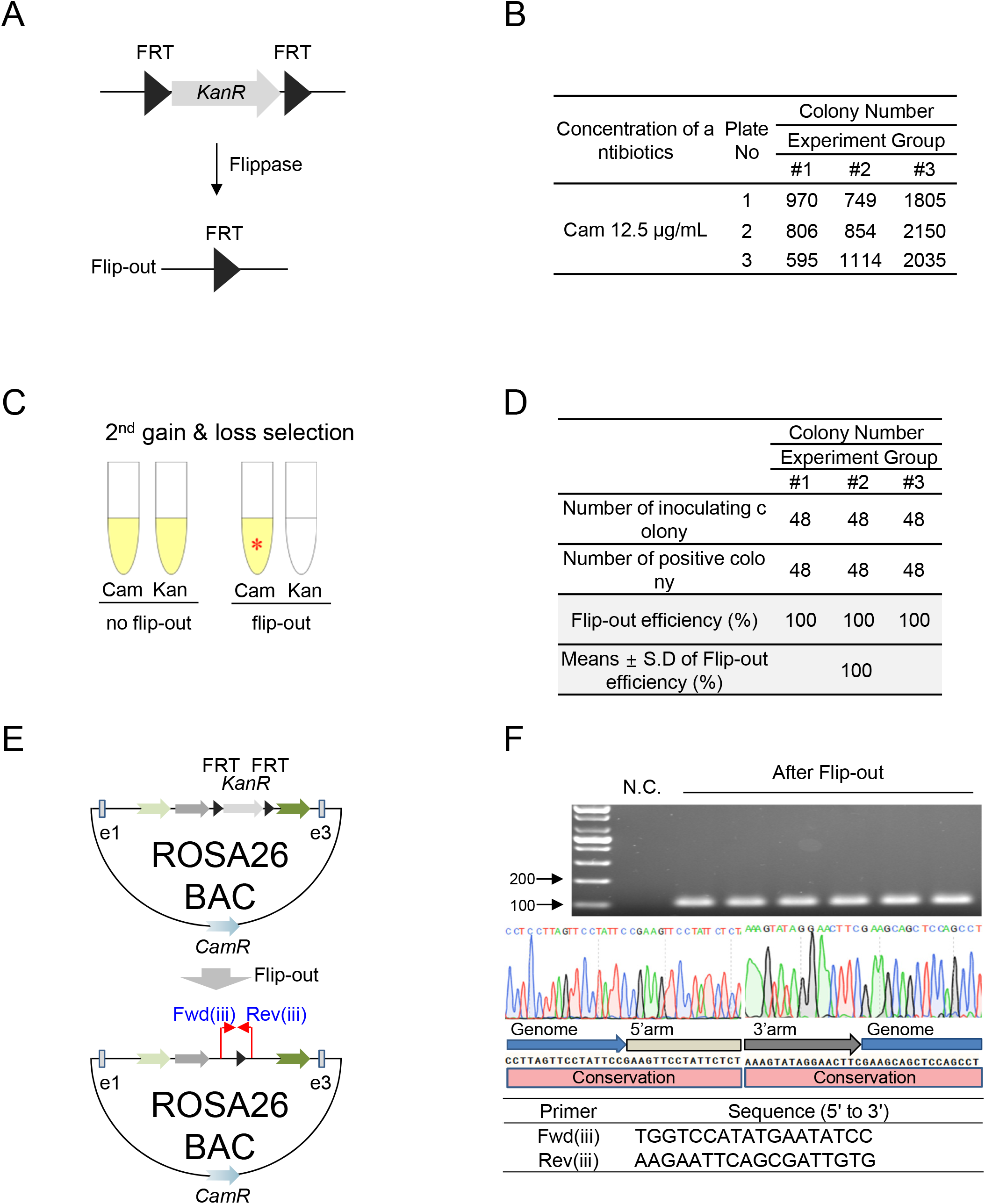
Detailed procedure and results for the flip-out reaction. (A) Overview of the flip-out reaction. In the presence of L-arabinose, the endogenous L-arabinose-inducible *FLP* gene in the SW105 strain will delete *KanR* flanked by two FRT sites. (B) Summary of colony numbers in LB plates containing Cam after the flip-out reaction. (C) Inoculation of positive clones into the 2^nd^ gain & loss selection system. (C) Illustration of the 2^nd^ gain & loss selection system for identifying recombinant clones. Red asterisk (*) indicates a flip-out positive clone. (D) Efficiency of flip-out reaction after performing the 2^nd^ gain & loss selection system. Means ± S.D., N = 3. (E) Flip-out scheme and location of colony PCR primers for confirming flip-out reaction. (F) Picture of agarose gel electrophoresis showing colony PCR results. PCR product was only amplified in the flip-out positive clone. Chromatogram results showing that the sequence of the amplified PCR product was conserved compared to the reference sequence at the border where flip-out was completed.

Following FLP induction, a LB plate containing Cam was used to discriminate the candidate clone(s). The number of candidate colonies appeared after 48 h incubation at 32°C was summarized in Fig. 2B. The 2^nd^ gain & loss screening system was used to determine whether the flip-out reaction was successful (Fig. 2C). The positive clones on the plate was inoculated and incubated for 24 h (Fig. 2C). This system will exhibit the two possible cases (Fig. 2C).

1. Case#1: positive in LB with Cam, positive with Kan (no flip-out)
2. Case#2: positive in LB with Cam, negative with Kan (flip-out)

Case#2 only occurs if the flip-out is successful. The average efficiency of obtaining a case#2 was 100% (Fig. 2D). To confirm the correct flip-out, colony PCR was conducted on all the clone from case#2. Two primers were located outside of the two FRT sites (Fig. 2E). The size of PCR product in flip-out region was approximately 100 bp, indicating the successful flip-out (Fig. 2F). Following PCR, DNA sequencing was performed to verify the flip-out region. All positive clones were found to exhibit recombinant sequences indicating 100% accuracy of 2^nd^ gain & loss screening system (Fig. 2F). Taken together, these results confirmed the accuracy of the 2^nd^ gain & loss screening system to 100%.

### 3^rd^ gain & loss screening system

To facilitate integration of the BAC construct into the genome, the Tol2 transposon system was used (Suster et al. 2011). The Tol2 transposon system yields the highest rate of genomic integration in the germ lineage resulting in the increased production of desired proteins (Suster et al. 2011; Balasubramanian et al. 2016). As *CamR* is located on a backbone vector of BAC clones, *CamR* was targeted to introduce the Tol2 transposon system. For the recombineering, a 5’ & 3’ *CamR* HR was introduced into the Cam targeting vector (CTV, Addgene ID: 131590) containing inverted left & right Tol2 transposons, *AmpR* for gain & loss screening, and a neomycin resistance (*NeoR*) gene for a mammalian selection marker (Fig. 3A). To purify chloramphenicol targeting cassette (CTC), CTV was cut with BamHI and XhoI (Fig. 3A).

**Fig. 3.**
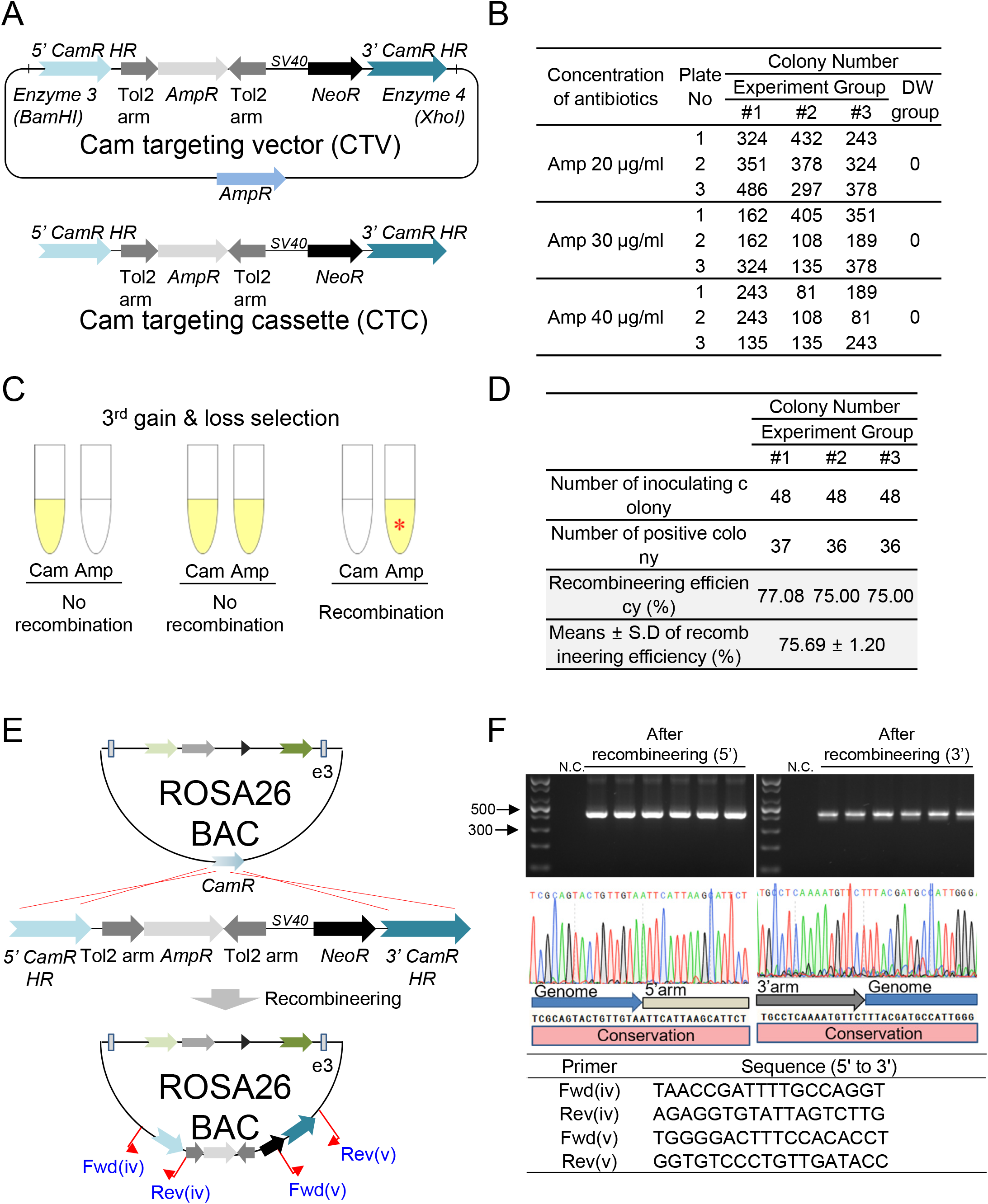
Detailed procedure and results for the 2^nd^ recombineering. (A) Overview of Cam targeting vector (CTV, Addgene ID: 131590) and Cam targeting cassette (CTC). (B) Summary of colony numbers in LB plates containing different concentrations of Amp after the 2^nd^ BAC recombineering. (C) Illustration of the 3^rd^ gain & loss selection system for identifying recombinant clones. Red asterisk (*) indicates the recombineering-positive clone. (D) Efficiency of the 2^nd^ recombineering after performing the 3^rd^ gain & loss selection system. Means ± S.D., N = 3. (E) 2^nd^ recombineering scheme and position of colony PCR primers for recombination confirmation. (F) Picture of agarose gel electrophoresis showing colony PCR results. PCR product was only amplified in the recombineering-positive clone. Chromatogram results showing that the sequence of the amplified PCR product was conserved compared to the reference sequence at the genome-CTC boundary.

As the recombination of CTC with a flip-out positive clone will lose *CamR* but acquire *AmpR* (from CTC), a LB plate containing Amp was used to discriminate the candidate clone(s). The number of candidate colonies appeared after 48 h incubation at 32°C was summarized in Fig. 3B. As CTV in supercoiled form cannot be efficiently cleaved with restriction enzymes, noncleaved CTVs may be included in the purified CTC. CTV contamination will result in a nonrecombinant clone, which has double antibiotic resistant genes, *AmpR* (from CTV) and *CamR* (from the BAC). To differentiate whether the clone derives from the desired recombination, the 3^rd^ gain & loss selection system was used. The candidate clones on the plate was inoculated in the 3^rd^ gain & loss selection system and incubated for 24 h (Fig. 3C). This system will exhibit the three possible cases (Fig. 3C).

1. Case#1: positive in LB with Cam, negative with Amp (no recombination)
2. Case#2: positive in LB with Cam, positive with Amp (no recombination)
3. Case#3: negative in LB with Cam, positive with Amp (recombination)

Case#3 only occurs if the recombineering is successful. The average efficiency of obtaining a case#3 was 75.69% (Fig. 3D). To confirm the correct recombination, colony PCR was conducted on all the clone from case#3. The Fwd primer was located in the BAC vector and the Rev primer in the CTC (Fig. 3E). If a recombineering occurred, the PCR product of 5’ recombineered and 3’ recombineered region should be approximately 400 bp and 390 bp, respectively (Fig. 3F). Following PCR, DNA sequencing was performed to verify the recombinant region. All positive clones were found to exhibit recombinant sequences indicating 100% accuracy of 3^rd^ gain & loss screening system (Fig. 3F). Taken together, these results also confirmed the accuracy of the third screening system to 100%.

### More strategies to increase recombination efficiency

The efficiency of the 2^nd^ BAC recombineering (75.69%) was 2 times higher than that of the 1^st^ BAC recombineering (31.94%). This discrepancy is presumed to be due to the difference in HR length, the decisive for the recombination efficiency (Sharan et al. 2009; Kung et al. 2013). However, the length of 5’ HR (473 bp) and 3’ HR (486 bp) in BTV was longer than 5’ HR (200 bp) and 3’ HR (200 bp) in CTV, so other factors may be involved in this discrepancy. Incomplete enzymatic digestion of the targeting vector increases the number of false positive clones, thereby reducing recombination efficiency (Jacobus and Gross 2015). Strategies to induce complete enzymatic digestion are required, but incomplete digestion may occur due to several other factors including DNA methylation (23). Thus, we hypothesized that strategy to distinguish between linear DNA and uncut circular DNA would increase the recombination efficiency. As a low percentage of agarose gel can distinguish fast-moving linear DNA fragments from slow-moving circular DNA (Lee et al. 2012), we used 0.2% agarose gel and differentiated the linearized BTC (4.7 kb; red rectangle) from uncut BTV (7.6 kb; yellow rectangle) (Fig. 4A). Then, the 1^st^ BAC recombineering was performed again. The efficiency of the 1^st^ BAC recombineering increased from 31.94% to 59.38%, indicating that a strategy to purify the targeting cassette on a low percentage of agarose gel can be another decisive factor for the efficient recombineering (Fig. 4A).

**Fig. 4.**
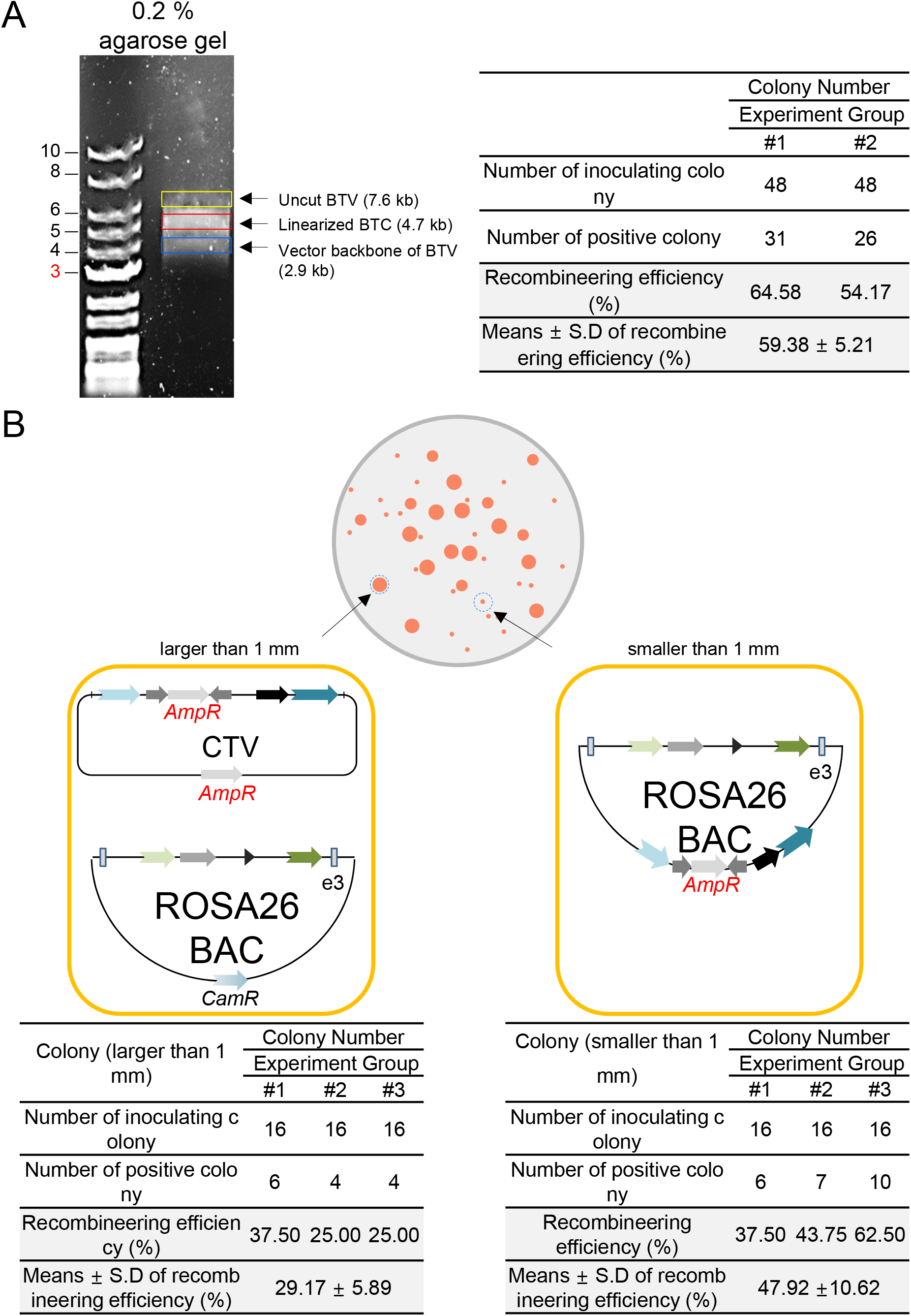
More strategies to increase recombination efficiency. (A) Picture of agarose gel electrophoresis showing the uncut BTV (7.6 kb; yellow rectangle), the linearized BTC (4.7 kb; red rectangle) and the vector backbone of BTV (2.9 kb; blue rectangle) after enzyme digestion of BTV with XhoI and XmaI. Efficiency of the 1^st^ recombineering after performing the 1^st^ gain & loss selection system. Means ± S.D., N = 2. (B) Illustration of colonies in LB plate containing Amp after the 2^nd^ BAC recombineering. After 48 h incubation at 32°C, colonies appeared. Large colonies (larger than 1 mm) were selected from one group and small colonies (smaller than 1 mm) from the other group. Efficiency of the 2^nd^ recombineering after performing the 3^rd^ gain & loss selection system in a larger or smaller colony-selected group. Means ± S.D., N = 3.

Targeting vector is high copy number plasmid, while BAC is low copy number plasmid. A high copy number plasmid replicates autonomously from the bacterial chromosome and is generally present in more than one copy per cell, providing higher antibiotic resistance (Jahn et al. 2016). CTV has *AmpR* for selection, so bacteria with uncut CTV with non-recombinant BACs can grow better than bacteria with recombinant BACs. Thus, we hypothesized that the selection of small colony will increase the recombination efficiency. Following the 2^nd^ recombineering, the candidate clones were distinguished using LB plates containing Amp. After 48 h incubation at 32°C, colonies appeared as shown graphically in Fig. 4B. To test our hypothesis, large colonies (larger than 1 mm) were selected from one group and small colonies (smaller than 1 mm) from the other group. The recombination efficiency obtained when selecting small colonies was increased by about 1.6-fold compared to when selecting large colonies (47.92% vs. 29.17%, Fig. 4B).

## DISSCUSSION

The rapid and efficient screening system is crucial for the BAC recombineering procedure. The importance of the screening system was realized by a strategy to reduce the number of false positive clones by performing long digestion of the targeting vector (Carreira-Rosario et al. 2013). However, due to many false positive clones, it still takes too much time to find a real recombinant clone (Carreira-Rosario et al. 2013). Another strategy have been attempted to avoid false positive clones from non-cleaved targeting vectors using PCR amplification of the targeting vector with PCR primers containing HR (Sharan et al. 2009). This strategy was also hampered by a false positive clones derived from PCR templates or unavoidable PCR errors during PCR amplification of the targeting vector (Sharan et al. 2009). In the current study, we uncovered a novel strategy in which simple inoculation of cells visually revealed the positive clones within 24 h. This strategy is based on whether cells can survive in the medium with specific antibiotics. Combination of survival or non-survival in a medium containing antibiotics was used a tool to determine whether a clone has a real recombined BAC or comes from a false positive background. The accuracy of this screening was confirmed by DNA sequencing results showing that all positive clones exhibited recombinant sequences. To our knowledge, our study provides the first demonstration that new gain & loss screening scheme enables 100% accuracy opening a new horizon in the field of BAC recombineering.

BAC recombination consists of time-consuming steps including 1^st^ recombineering, flip out and 2^nd^ recombineering. This time-consuming procedure has been a major obstacle for researchers starting BAC recombineering. Procedures to reduce BAC recombination duration are the most demanding criteria for researchers to conduct BAC recombination. The 100% accurate gain & loss screening system facilitated recombineering process by making each recombination process successful at once. Furthermore, the increased recombination efficiency by using long HR promoted recombination progress by allowing identification of the recombinant clones out of only 48 colonies per step. The efficiency of the 1^st^ and 2^nd^ BAC recombineering was 31.94% and 75.69%, respectively, whereas that of traditional BAC recombineering with short HRs was less than 10% (Liu et al. 2003). Thus, our new strategy allowed us to complete the entire BAC recombineering procedure within just 13 days (Fig. 5). Taken together, considering short duration, we expect that this protocol will open a new path for laboratories with less experience in the field of BAC recombineering.

**Fig. 5.**
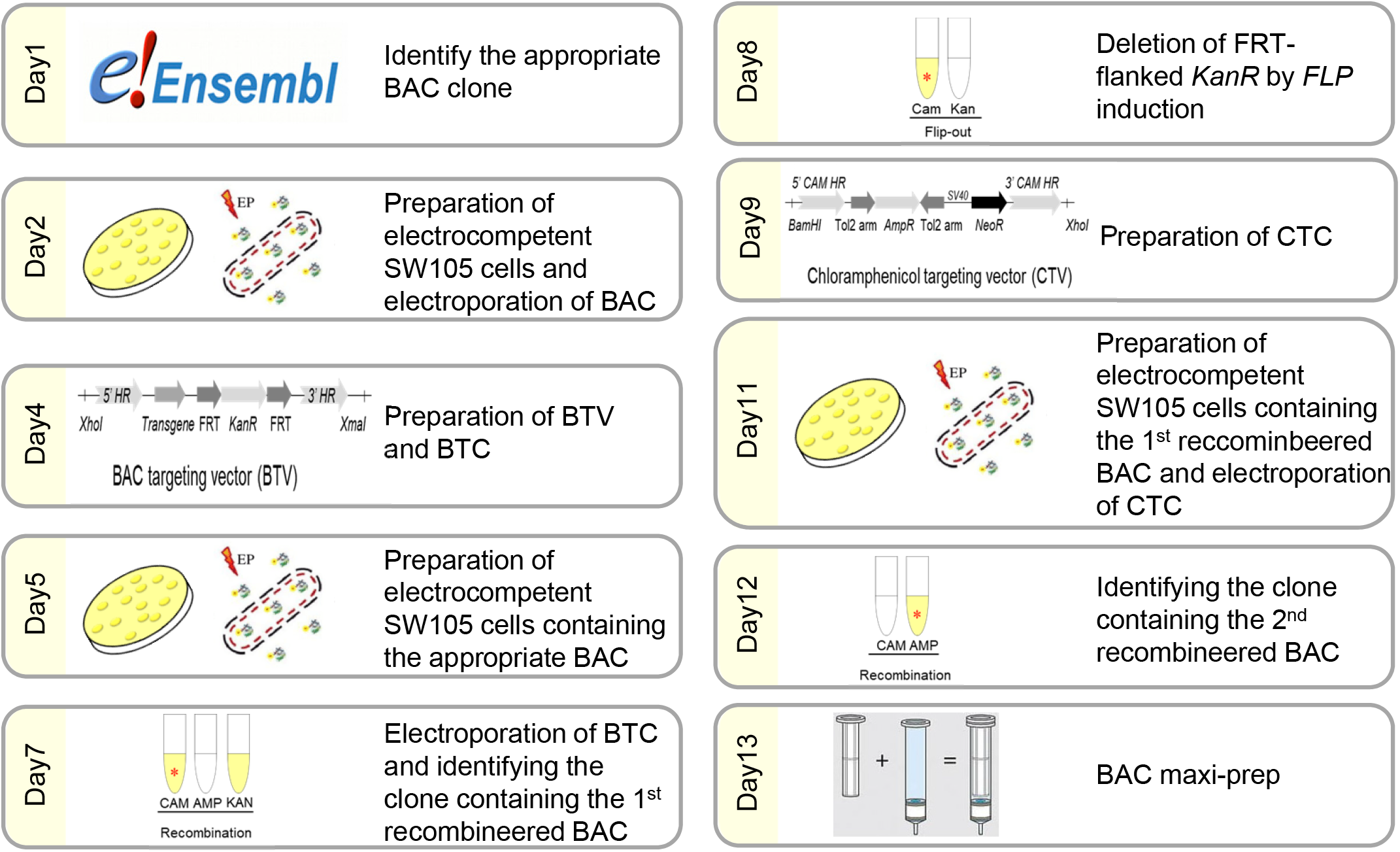
Summary of the experimental procedure and timing for each step.

Recombineering is a genetic and molecular biology technique based on homologous recombination systems, which mediate efficient recombination of linear DNA molecules flanked by long HR (Thomason et al. 2014). The significance of long HR is highlighted by the finding that the targeting cassette with long HR (200–500 bp) is more efficient for recombineering than the targeting cassette with short HR (approximately 50 bp) (Degryse 1996; Sharan et al. 2009; Dickinson et al. 2015). Targeting vectors with long HR were linearized by restriction digestion and purified in preparation for subsequent use in BAC recombineering steps. Long-term DNA digestion is crucial for complete digestion, but incomplete digestion can occur if enzyme activity is blocked by DNA methylation (Snounou and Malcolm 1984). As non-cleaved targeting vectors are inevitable, a new strategy is needed to avoid false positive clones. Here, we found that a novel tool using a low percentage of agarose gel increased the likelihood of obtaining recombinant clones by effectively avoiding uncut targeting vectors. The significance of a novel approach was supported by the results showing that the 1^st^ BAC recombination efficiency increased from 31.94% to 59.38%. Taken together, our results imply that this new approach can be considered an alternative to solving problems that may arise when using long HR in targeting vectors.

BAC is based on an F-factor plasmid that maintains a small number of copies in bacterial cells, while the targeting vector is a high copy number plasmid (Asami et al. 2011). A low copy number plasmid has only one or a few copies in each bacterium, so antibiotic resistance is low compared to high copy number plasmids (Jahn et al. 2016). Thus, bacteria with low copy number plasmid grow slower than bacteria with high copy number plasmid in the presence of antibiotics (Trivedi et al. 2014). In this study, we proposed a novel strategy that use the growth differences between bacteria with recombinant BAC and bacteria without recombinant BAC. Bacteria with recombinant BAC are presumed to grow slower than bacteria with both unrecombined BAC and targeting vectors. This strategy was validated by the results showing that the recombination efficiency obtained when selecting small colonies was 1.6 times higher than when selecting large colonies. Based on these findings, we conclude that a strategy using growth differences in the presence of antibiotics will be another alternative that promotes fast and efficient BAC recombination.

In summary, we have developed a new strategy that makes it easy to visually confirm the success of BAC recombineering and reduce the overall processing time to less than two weeks (13 days) (Fig. 5). This new screening system will provide significant advances in current methodologies for BAC recombineering and might be of broad interest to many researchers in different fields including biopharmaceutical protein expression and other large-vector applications for various transgenic animals.

## Author Contributions

MUK and JTP conceived of and designed the experiments. MUK performed the experiments. MUK, SO, and JTP wrote and edited the paper.

## Acknowledgements

This research was supported by a Basic Science Research Program through the National Research Foundation of Korea (NRF) funded by the Ministry of Science, ICT, and Future Planning (NRF-2018R1D1A1B07040293). This research was also supported by Research Assistance Program (2019) in the Incheon National University.

## Financial disclosure

The authors declare no conflicts of interest.

## Information pertaining to writing assistance

The authors declare no writing assistance

## Ethical disclosure

The authors state that they have followed the principles outlined in the Declaration of Helsinki for all human or animal experimental investigations. In addition, for investigations involving human subjects, informed consent has been obtained from the participants involved.

## Data sharing statement

The authors declare that they shall make data available to the scientific community without any restrictions.

---

**Supplementary Fig. 1.**
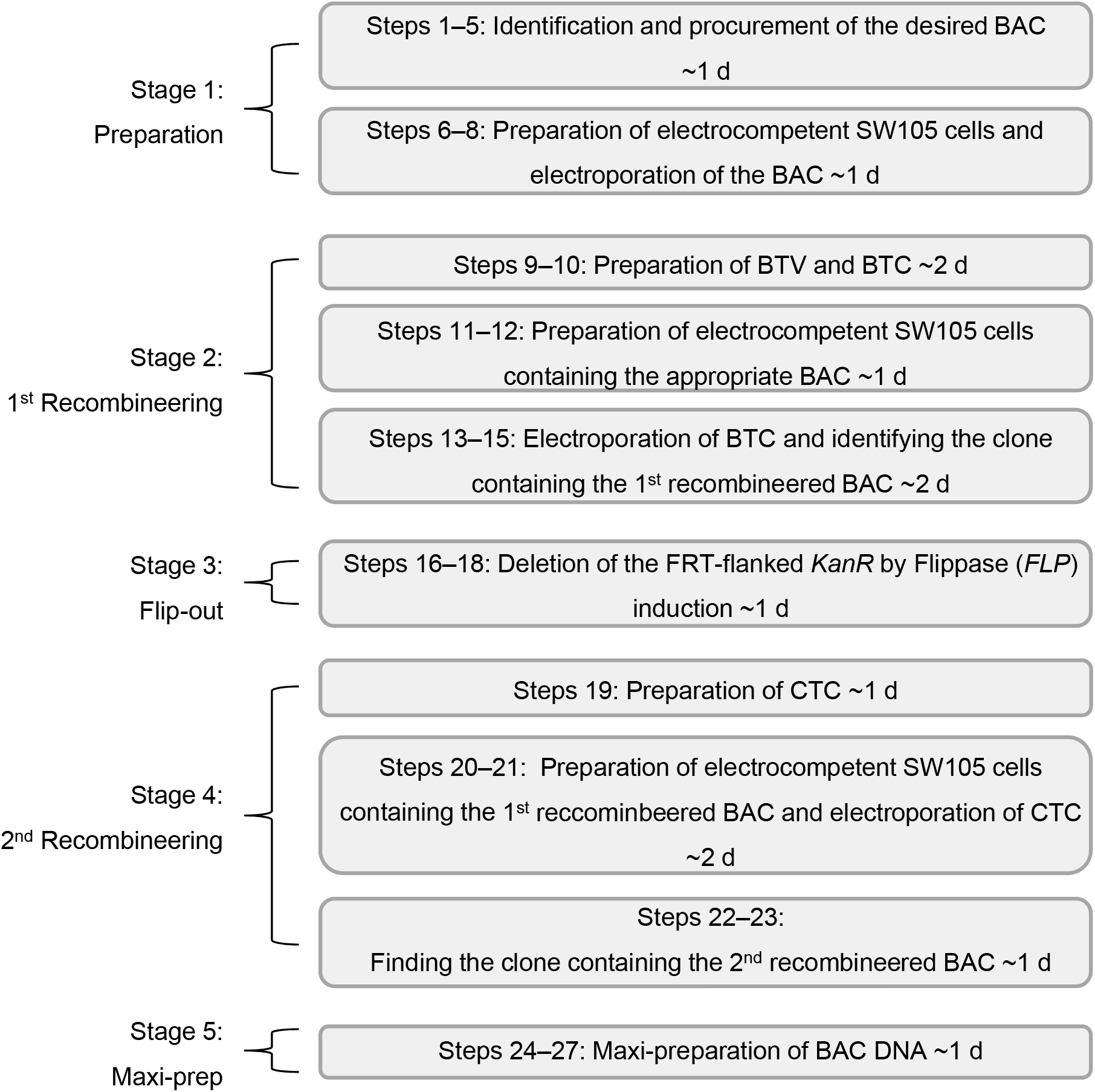
Overview of the experiment procedure and estimated timing of each step.

**Supplementary Fig. 2.**
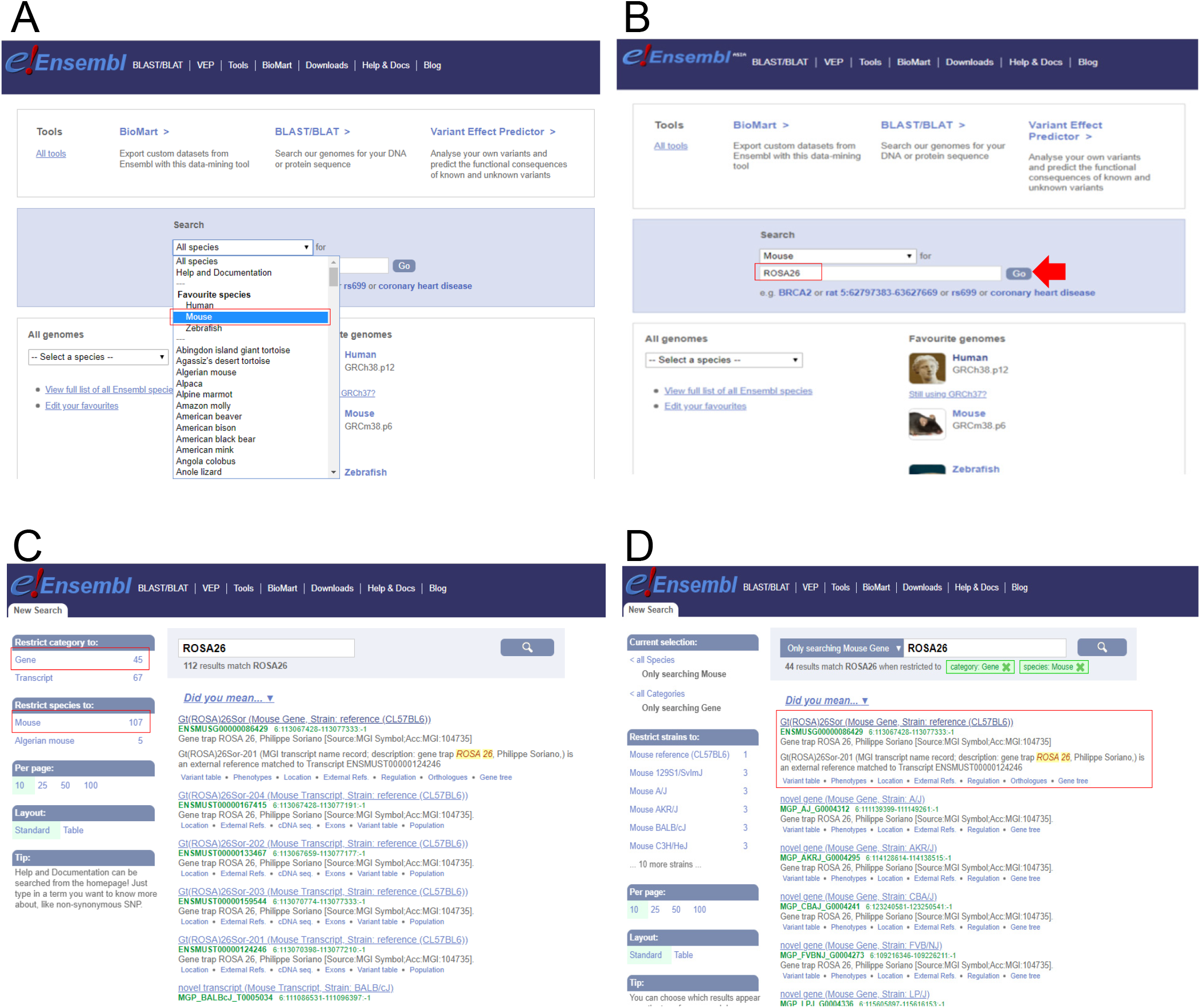
Identification and procurement of the desired BAC.

**Supplementary Fig. 3.**
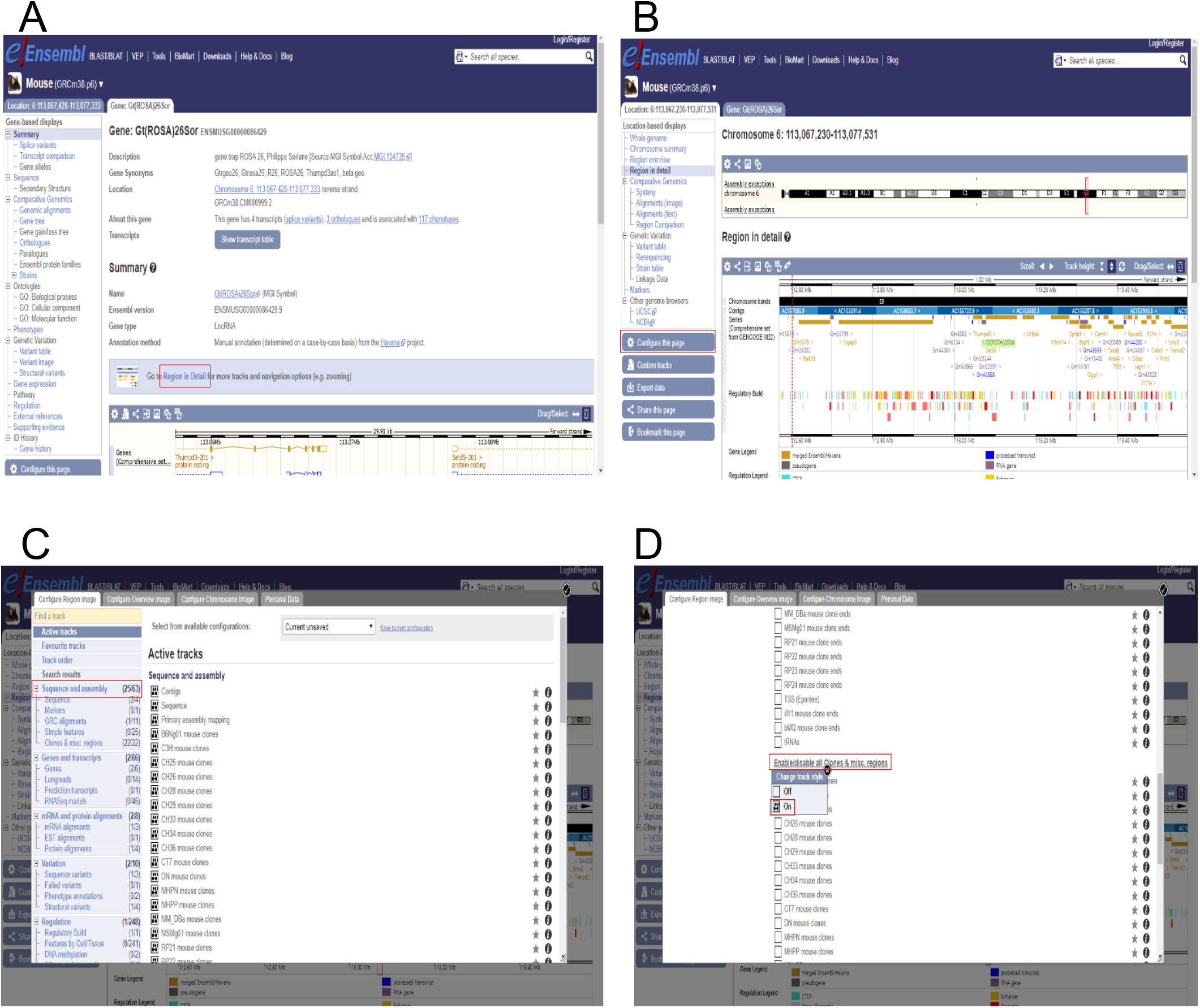
Identification and procurement of the desired BAC.

**Supplementary Fig. 4.**
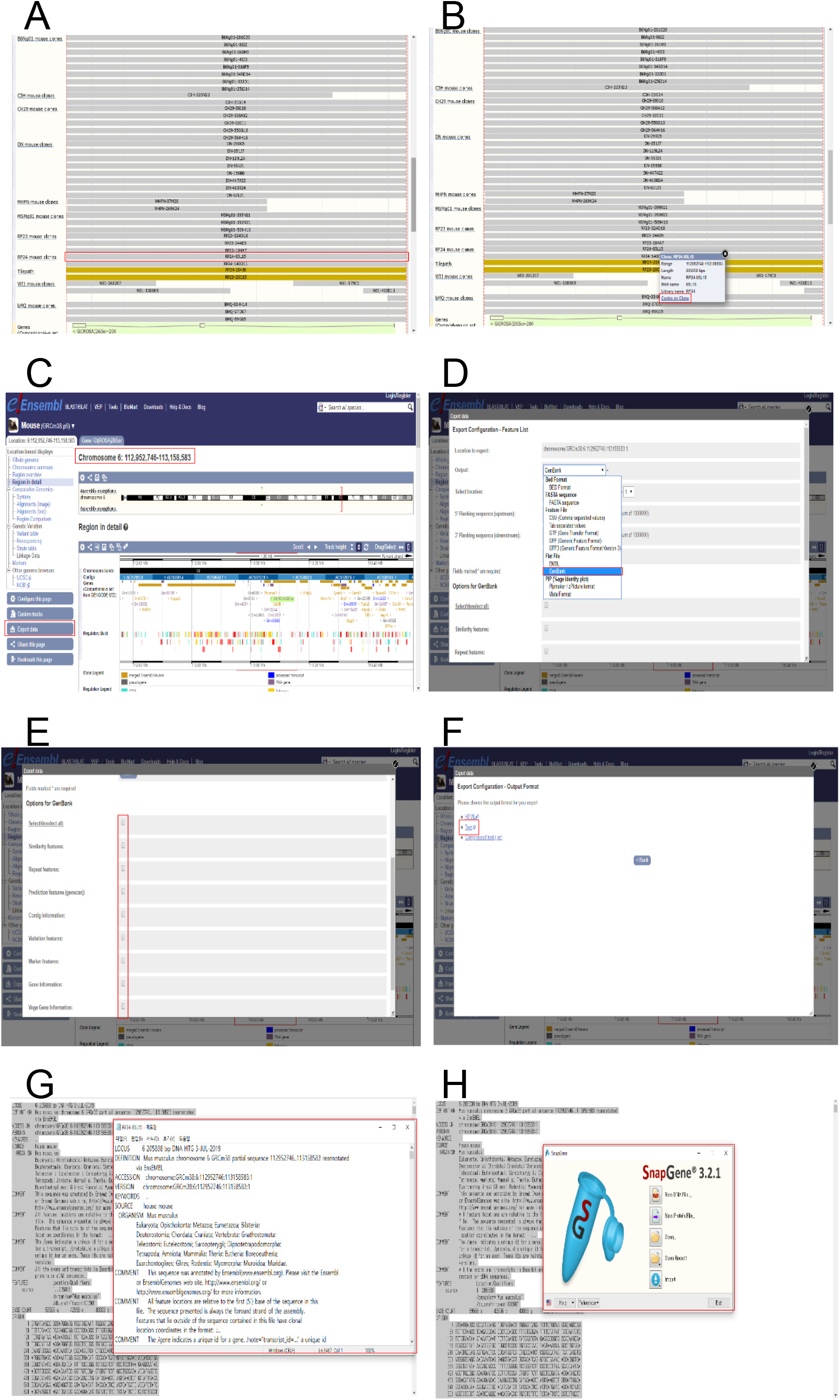
Identification and procurement of the desired BAC.

**Supplementary Fig. 5.**
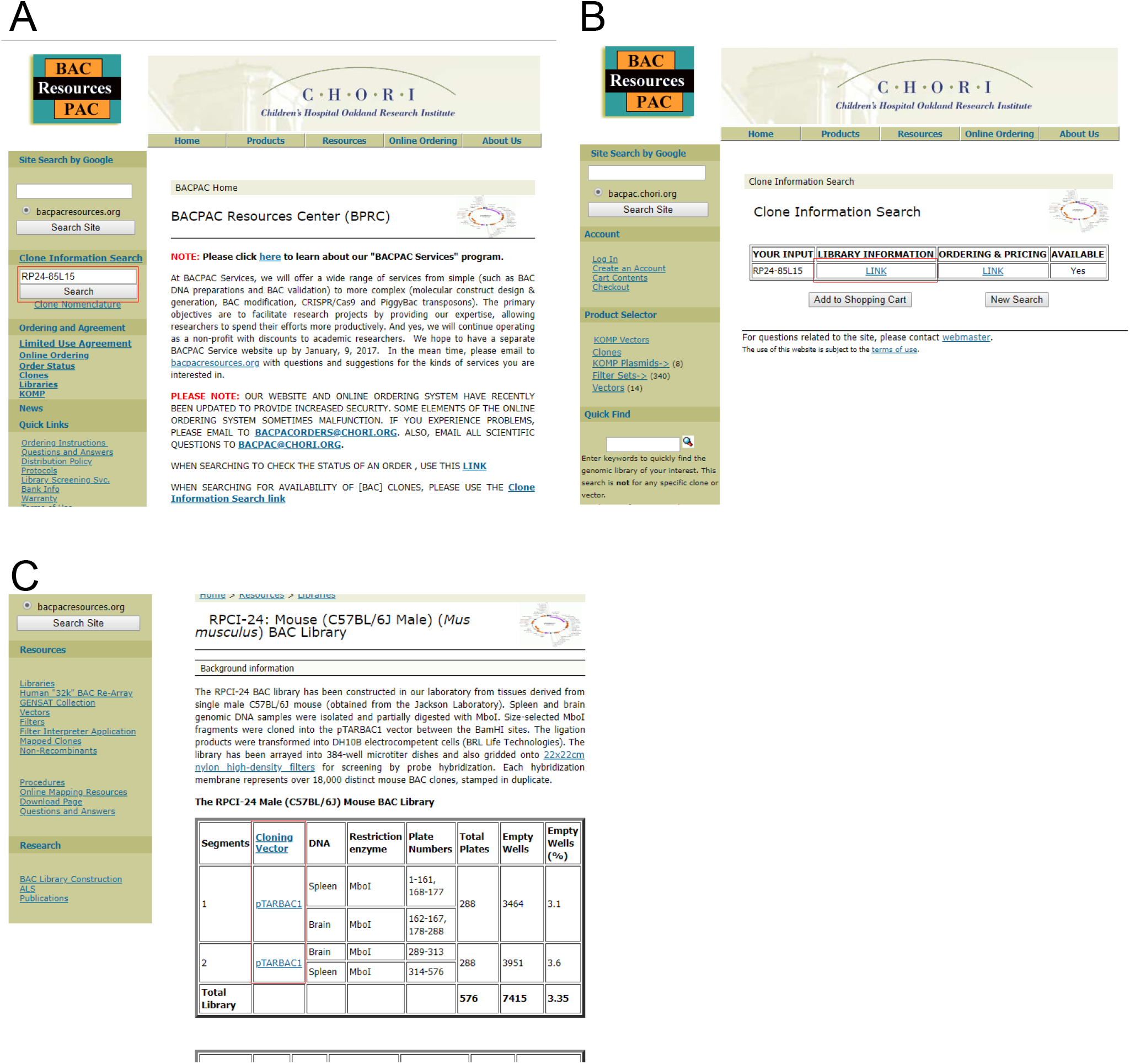
Identification and procurement of the desired BAC.

**Supplementary Fig. 6.**
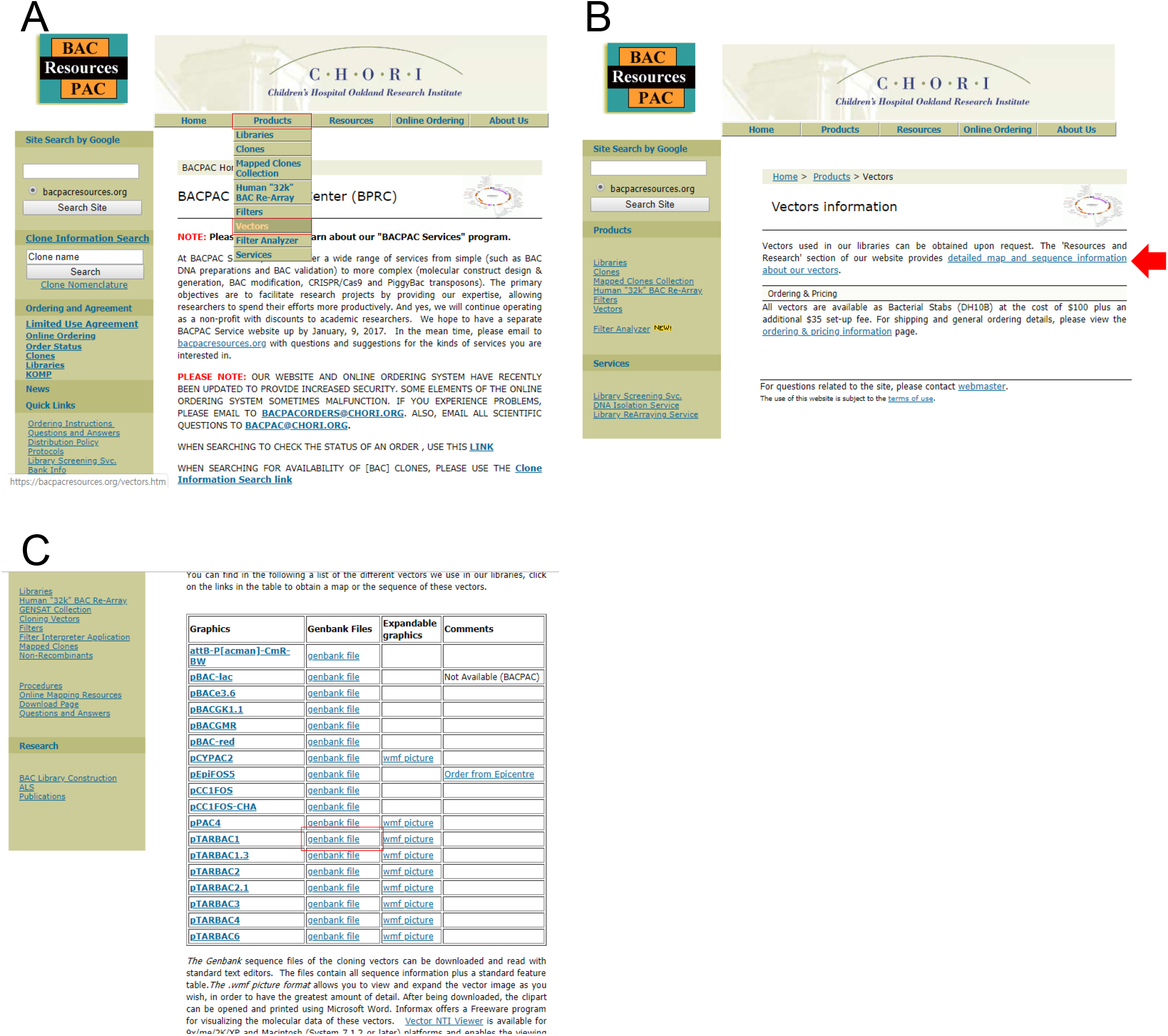
Identification and procurement of the desired BAC.

